# A qPCR assay for *Bordetella pertussis* cells that enumerates both live and dead bacteria

**DOI:** 10.1101/782177

**Authors:** Stacy Ramkissoon, Iain MacArthur, Muktar Ibrahim, Hans de Graaf, Robert C. Read, Andrew Preston

## Abstract

*Bordetella pertussis* is the causative agent of whooping cough, commonly referred to as pertussis. Although the incidence of pertussis was reduced through vaccination, during the last thirty years it has returned to high levels in a number of countries. This resurgence has been linked to the switch from the use of whole-cell to acellular vaccines. Protection afforded by acellular vaccines appears to be short-lived compared to that afforded by whole cell vaccines. In order to inform future vaccine improvement by identifying immune correlates of protection, a human challenge model of *B. pertussis* colonisation has been developed. Accurate measurement of colonisation status in this model has required development of a qPCR-based assay to enumerate *B. pertussis* in samples that distinguishes between viable and dead bacteria. Here we report the development of this assay and its performance in the quantification of *B. pertussis* from human challenge model samples. This assay has future utility in diagnostic labs and in research where a quantitative measure of both *B. pertussis* number and viability is required.

## Introduction

Whooping cough, or pertussis, is a highly contagious respiratory tract infection of humans caused by the gram-negative coccobacillus *Bordetella pertussis*. Clinical manifestations of pertussis depend on age and immune status of the host and include a low-grade fever, cyanosis, and paroxysmal cough accompanied by a high-pitched “whoop” (1). Infants aged less than 1 year old present the highest incidence of pertussis and are also at the greatest risk of severe disease and death (2). The introduction of vaccination in the early 1950s significantly reduced the incidence of pertussis in developed nations, however the number of reports of pertussis has been progressively increasing over the last thirty years (3). For example, in the UK, Public Health England has reported a greater than ten-fold increase in pertussis cases over the eight-year period of 2005-2013 (4). This rise has been echoed in other countries including Australia, the Netherlands, and the US (5–10).

The reason for this resurgence is not certain, however it has been strongly linked to the switch from using whole-cell vaccines (WCVs) to using acellular vaccines (ACVs). ACV-induced immunity appears to wane more quickly than WCV-induced immunity. In baboons, compared to WCV-induced immunity, ACV-induced immunity protects from disease, but does not prevent colonization by *B. pertussis* or prevent transmission of the bacteria to other hosts (11–12). In addition, in many countries using ACVs, there has been a dramatic increase in the isolation of *B*. pertussis deficient for the production of the ACV-vaccine antigen pertactin. In ACV-immunised hosts pertactin-deficient *B. pertussis* may have a fitness advantage over pertactin-producing isolates, raising concern that the use of ACVs is selecting for vaccine escape strains of *B. pertussis* (13). These issues have highlighted the need to better understand the detailed differences between WCV and ACV induced immune responses and the immune response to infection, and to identify biomarkers of protective immunity to *B. pertussis* infection. This would aid the evaluation of the efficacy of future *B. pertussis* vaccines that might be needed to combat *B. pertussis* resurgence. To this end, a human challenge model of *B. pertussis* colonisation has been developed as part of the EU-funded PERISCOPE Project (14–15). In this model it is necessary to be able to monitor the colonisation status of participants at frequent intervals. Current detection methods for *B. pertussis* include culture from nasopharyngeal swabs or other nasopharyngeal samples. However, *B. pertussis* is slow-growing and takes several days to produce visible growth on laboratory agar. A more rapid method would improve safety for human challenge model volunteers. Real-time PCR detection (qPCR) of *B. pertussis* DNA provides rapid identification of *B. pertussis* within hours and for diagnosis of *B. pertussis* infection is regarded as more sensitive than culture. However, traditional qPCR assays cannot distinguish between viable and dead bacteria, which is essential to determine whether participants are actively colonised.

Here we report the modification of a standard qPCR assay used for laboratory diagnosis of *B. pertussis*, through treatment of samples with propidium monoazide (PMA) that inhibits PCR-mediated amplification of DNA from dead cells and allows distinguishing of viable from dead cells (16–20). The use of PMA involves an initial incubation of samples with PMA in darkness, during which it diffuses into dead cells, followed by light activation of PMA that permanently modifies the gDNA of dead cells, preventing it from acting as a template in PCR. The optimisation of this assay and its use to enumerate viable and dead *B. pertussis* from human challenge model samples is described. In addition, this assay has wider uses in diagnostic and other research settings where a quantitative measure of viable *B. pertussis* number is required.

## Materials and methods

### Bacterial strains and culture conditions

*B. pertussis* strain BP1917 is a wild-type strain considered representative of currently circulating *B. pertussis* (21). It was cultured on charcoal agar at 37°C for 3 days for routine culture.

### The preparation of heat-killed bacterial cell suspensions

Plate-grown B1917 were resuspended in PBS to an OD_600_ = 1.0 (approximately 10^9^ cfu/ml). To optimise heat killing, 1 ml aliquots were heat-killed at 80°C for 1, 3 and 6 minutes in a pre-heated heat block. Aliquots were placed on ice immediately after incubation. Bacterial death was confirmed by the absence of growth after streaking 10 μl of suspension onto charcoal agar plates and incubating at 37°C for 5 days. To ascertain the integrity of heat-killed cells, samples were subjected to flow cytometry (FACSCantoII, BD UK Ltd, Wokingham, U.K.). A detergent-lysed sample acted as a positive control for lysis and a sample containing live cells was a positive control for cell integrity.

### The preparation of THP-1 cells

THP-1 (ATCC® TIB-202™) cells were maintained in RPMI 1640 medium, fetal bovine serum (10%), 1% streptomycin, penicillin and glutamine (ThermoFisher Scientific, Loughborough, UK) as per standard methods. Heat-killed THP-1 cells were prepared by incubating cell suspensions at 10^5^ cells/ml at 80°C for 6 minutes in a pre-heated heat block.

### Optimisation of PMA treatment conditions

PMA Dye, 20 mM in H_2_O (Cambridge BioSciences, Cambridge, UK), was stored at −20°C in the dark, thawed on ice and added to 2 ml clear centrifuge tubes containing 200 μl of cell suspensions to a final concentration of 20 μM, 30 μM, or 50 μM. PMA-free samples served as controls for each condition tested. Tubes were covered with aluminium foil and shaken on an orbital shaker for 5, 10, 20, 30 or 70 minutes. Samples were then exposed to light using the PMA-Lite LED Photolysis Device (Cambridge BioSciences, Cambridge, UK) for 5, 10, 20, 30 or 40 minutes. Samples were pelleted using the Heraeus Pico 17 Centrifuge at 2000xg (ThermoFisher Scientific, Loughborough, UK) for 10 minutes at room temperature prior to DNA isolation.

### Genomic DNA Isolation

Genomic DNA (gDNA) was isolated using the GenElute Bacteria Genomic DNA Kit (Sigma-Aldrich, Dorset, UK) according to the manufacturer’s instructions and eluted with 200 μl of elution buffer. gDNA was purified from THP-1 cells using the QIAmp DNA mini and blood extraction kit (QIAgen, Manchester, UK) as per the manufacturer’s protocol. gDNA was quantified using a Qubit 1.0 fluorometer (Invitrogen, Loughborough, UK) according to the manufacturer’s instructions.

### Quantitative PCR

qPCR was performed using Sybr green PCR Master Mix (Applied Biosystems, Loughborough, UK). The final reaction volume was 25 μl comprising of 12.5 μl Sybr Green master mix, 7.3 μl H_2_O, 0.1 μl of 100 nM stocks of each primer, and 5 μl of template sample. The reaction was run using a StepOne Plus RT PCR System (Applied Biosystem, Loughborough, United Kingdom) with the cycle conditions described in Table 1.

**Table 1.** Thermocycling Conditions for qPCR using Sybr Green:

| STEP | TEMP | TIME |
| --- | --- | --- |
| Initial Denaturation | 95°C | 10 minutes |
| 40 Cycles | 95°C | 15 s |
|  | 48°C | 60 s |
|  | 60°C | 60 s |
| Melt Curve Analysis | 95°C | 15 s |
|  | 48°C | 60 s |
|  | 60°C | 60 s |

Alternatively, qPCR was performed using a fluorogenic probe (Eurofins, Ebersberg, Germany). The reaction volume was 20 μl comprising of 2 μl of 1x Taqman Gene Expression Mastermix (Applied Biosystems, Loughborough, UK), 2 μl of 900 nM stocks of each primer, 2 μl of 150 nM stock of probe, 2 μl of nuclease-free water and 2 μl of template sample. The reactions were run using the StepOne Plus RT PCR System using the cycling parameters found in Table 2. The sequence of primers and probe were as described previously (22): forward primer (5’ATCAAGCACCGCTTTACCC 3’), reverse primer (5’ TTGGGAGTTCTGGTAGGTGTG 3’) and probe (5’ AATGGCAAGGCCGAACGCTTCA 3’) was labelled with FAM and Black Hole Quencher.

**Table 2.** TaqMan Thermocycling Conditions for qPCR:

| STEP | TEMP | TIME |
| --- | --- | --- |
| Step 1 |  |  |
| Holding Stage | 50°C | 2 minutes |
| Step 2 | 95°C | 10 minutes |
| 40 Cycles | 95°C | 15 s |
|  | 60°C | 60 s |

### Calculating copy number from Ct values/ DNA Concentration

The genome copy number equivalent to the amount of template in a qPCR reaction was calculated using the formula:

copy number = (amount of template in ng * 6.022×10^23^) / (length of genome in bp × 1×10^9^ * 650). The genome of BP1917 is 4,102,186 bp (21).

### Preparation of bacterial and THP-1 cell suspensions

To evaluate if eukaryotic cells interfere with the enumeration of live *B. pertussis* cells using qPCR, 10^3^ live *B. pertussis* were combined with THP-1 gDNA equivalent to 10843, 8414, 5385, 3446, 2804, 2316, 1868, 1503, 1251, 1023, 875, 746, 671, 507, 366, or 275, 141, 29 cells. A sample without THP-1 DNA served as a control.

To evaluate the possible sequestration of PMA by eukaryotic DNA, 10^6^ heat-killed *B. pertussis* were combined with either 100,000 heat-killed THP-1 cells, 100,000 live THP-1 cells or without THP-1 cells and were then treated with the selected PMA treatment. Non PMA-treated samples were run in parallel.

To determine if eukaryotic cells interfered with the action of PMA on dead bacterial cells, 100,000 live THP-1 cells were combined with different ratios of viable *B. pertussis* cells and heat-killed *B. pertussis* cells (final bacterial concentration was 10^6^ cfu/ml) in a clear Eppendorf tube, total volume 200 μl. These samples were then subjected to the selected PMA treatment. A non-PMA treated control was included. gDNA was extracted from each sample and used for qPCR.

### Statistical Analysis

Unpaired T tests, corrected for multiple comparisons, and two-way ANOVA using the Holm-Sidak method were used to evaluate statistical significance. One-way ANOVA and Dunnett’s multiple comparisons test, with a single pooled variance was also used. A p value of <0.05 was defined as statistically significant and is indicated by asterisks.

## Results

### qPCR provides a lower limit of detection of 2 *B. pertussis* cells

IS481 is often used as the target for qPCR detection of *B. pertussis* as it is present at ~250 copies per cell in *B. pertussis*, providing great sensitivity. To develop a PMA-qPCR assay, the sensitivity of qPCR for detection of *B. pertussis* was tested over a range of template gDNA concentrations. A linear relationship between Ct value and template concentration was observed over the range of 2 to approximately 2.42×10^6^ B1917 cells for qPCR (Figure 1). Ct values greater than 35 were considered to be a negative reaction. Probe based detection was more sensitive than sybr-green based detection (data not shown). Thus, the assay is able to detect *B. pertussis* gDNA equivalent to very few bacterial cells and is linear over a wide range of *B. pertussis* concentrations.

**Figure 1.**
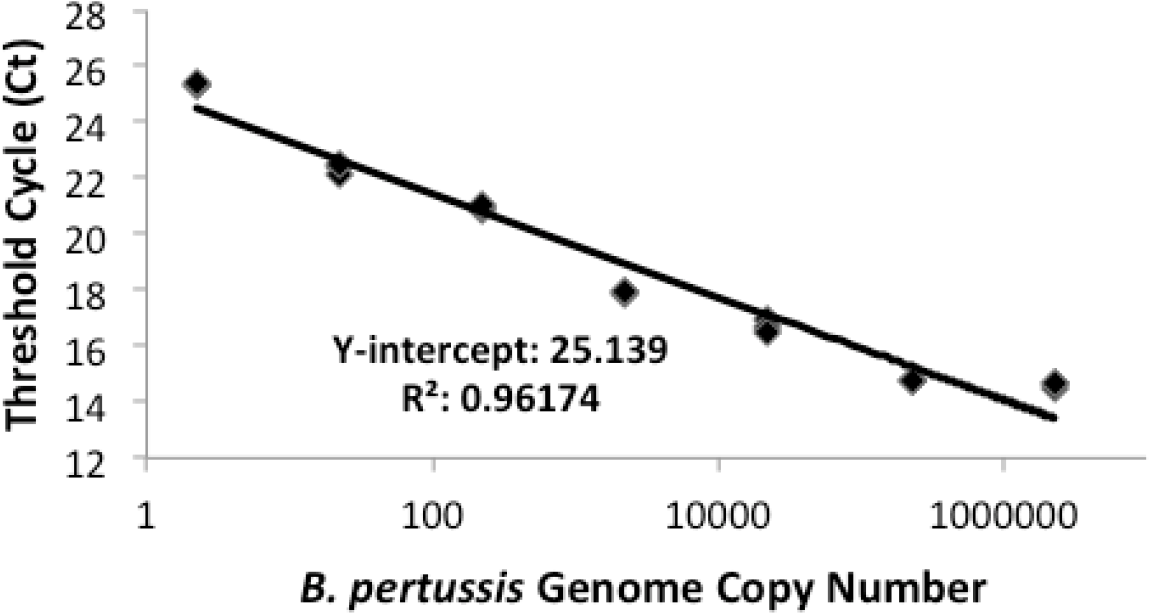
Standard curve of Ct value versus template concentration. Template DNA concentration is expressed as B1917 genome copy number. The linearity was determined to be from 2 to approximately 2.42×10^6^ B1917 genomes for qPCR.

### Heat-killing B. pertussis at 80°C for 6 minutes maintained the integrity of cells

The ability of PMA to inhibit PCR-amplification from dead *B. pertussis* was tested using heat-killed *B. pertussis*. It was envisaged that clinical samples may contain dead, but intact, *B. pertussis*. Heat-killing may cause cell lysis which would not mimic intact dead cells. Thus, the integrity of cells following heat killing was assessed using flow cytometry. Incubation of *B. pertussis* suspensions at 80°C for 6 minutes resulted in 100% killing, but with cells remaining intact and were the conditions used throughout (Figure 2).

**Figure 2.**
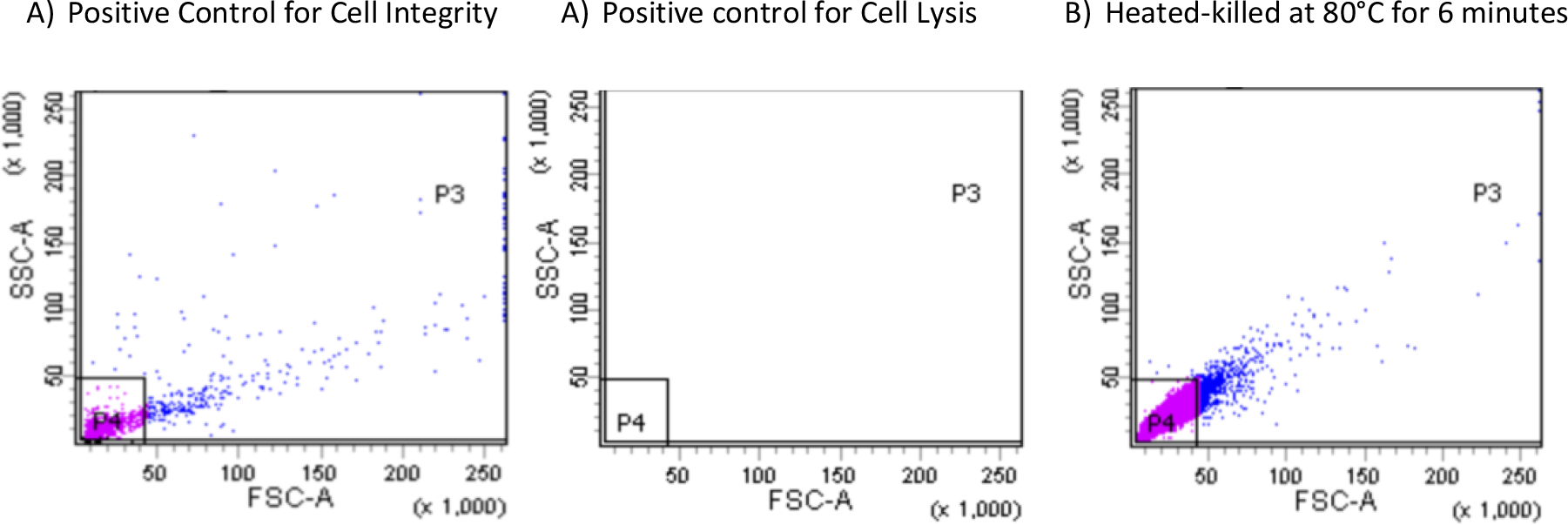
The effect of heat killing on the integrity of *B. pertussis* cells, measured by flow cytometry. A) Positive control for cell integrity – a suspension of live *B. pertussis*; B) Positive control for cell lysis – detergent lysed *B. pertussis*; C) Heat-killed *B. pertussis* suspension. The heat-killed *B. pertussis* suspension incubated for 6 minutes at 80°C displayed similar scatter as the live cell suspension. No particles were detected in a suspension of detergent-lysed *B. pertussis*. Therefore, cells remained intact in the heat-killed *B. pertussis* suspension incubated for 6 minutes at 80°C when compared to the positive cell integrity control.

### Optimisation of PMA treatment

The effect of PMA concentration on inhibition of PCR amplification from dead *B. pertussis* was tested (Figure 3). Incubation of heat-killed cells with 50μM of PMA resulted in a 97.42 % reduction in PCR signal compared to that generated from untreated samples. Lower levels of PMA also resulted in very similar levels of inhibition (Figure 3).

**Figure 3.**
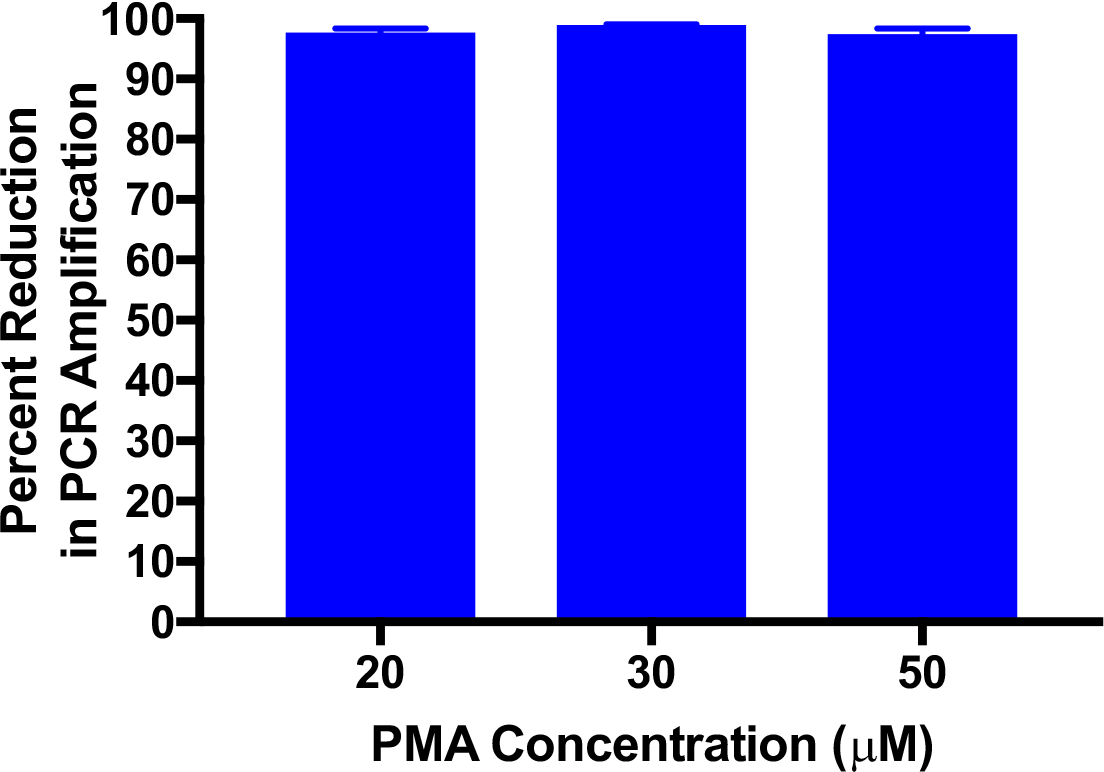
The effect of PMA concentration on the reduction of the PCR amplification signal from heat-killed cells. Treatment of samples with either 20 μM, 30 μM, or 50 μM of PMA produced a ≥97% reduction in the PCR amplification signal compared to untreated samples. Error bars represent standard deviations from two biological replicates. Data from a representative experiment repeated three times.

The optimal conditions for photo-activation of PMA were determined. Incubation under dark conditions for 10 minutes followed by light activation for between 5 and 30 minutes resulted in greater than 99% reduction in PCR signal from dead cells compared to untreated controls. Five minutes of light activation following 10 minutes of darkness resulted in 99.64% reduction in detection of *B. pertussis* DNA (Figure 4).

**Figure 4.**
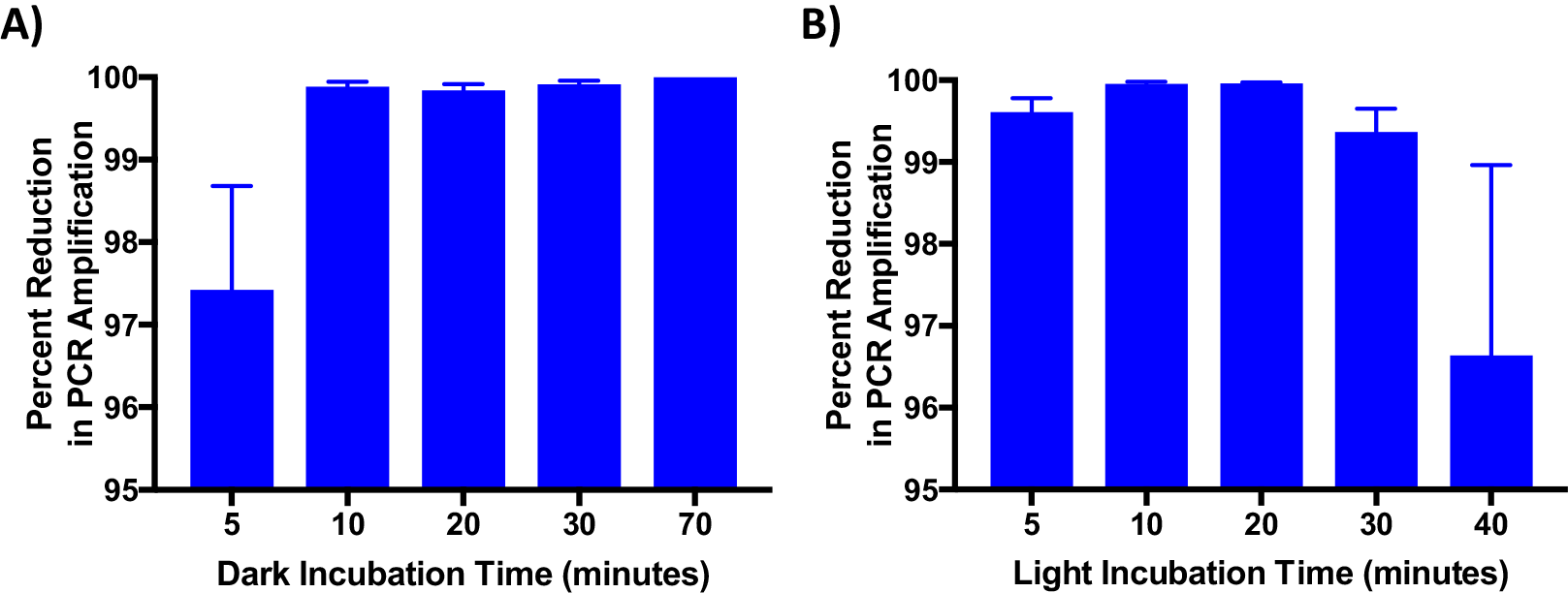
The effect of dark and light exposure times on PMA-inhibition of PCR amplification. A) Dark incubation B) Light Incubation. PMA and untreated heat-killed suspensions were incubated for 10, 20, 30, and 70 minutes in the dark followed by exposure to 5 minutes of light. 10 minutes or longer of incubation in the dark produced a ≥99% reduction in the PCR amplification signal. Optimal light incubation periods were determined by incubating untreated and PMA treated heat-killed suspensions in the dark for 10 minutes followed by light exposure for 5, 10, 20, 30, and 40 minutes. Incubating PMA treated heat-killed samples under light for periods of 5, 10, 20, and 30 minutes produced a 99% or greater reduction in the PCR amplification signal. Five minutes was selected as the standard light incubation period. Error bars represent standard deviation from two biological replicates. The experiment was repeated with the same result.

From these optimisations, standard conditions of 50uM PMA and incubation in the dark for 10 minutes followed by light activation for 5 minutes were selected as minimal incubation times that achieved high levels of inhibition. Even though 20uM PMA inhibited PCR amplification from dead cells, 50uM PMA was selected as the concentration to use in the assay, as clinical samples will contain cells other than *B. pertussis* that may sequester PMA, requiring an excess for consistent inhibition of PCR signal from dead *B. pertussis*. These conditions were tested in four independent assays. An average of 94.15% reduction in PCR signal was observed compared to untreated controls (Figure 5).

**Figure 5.**
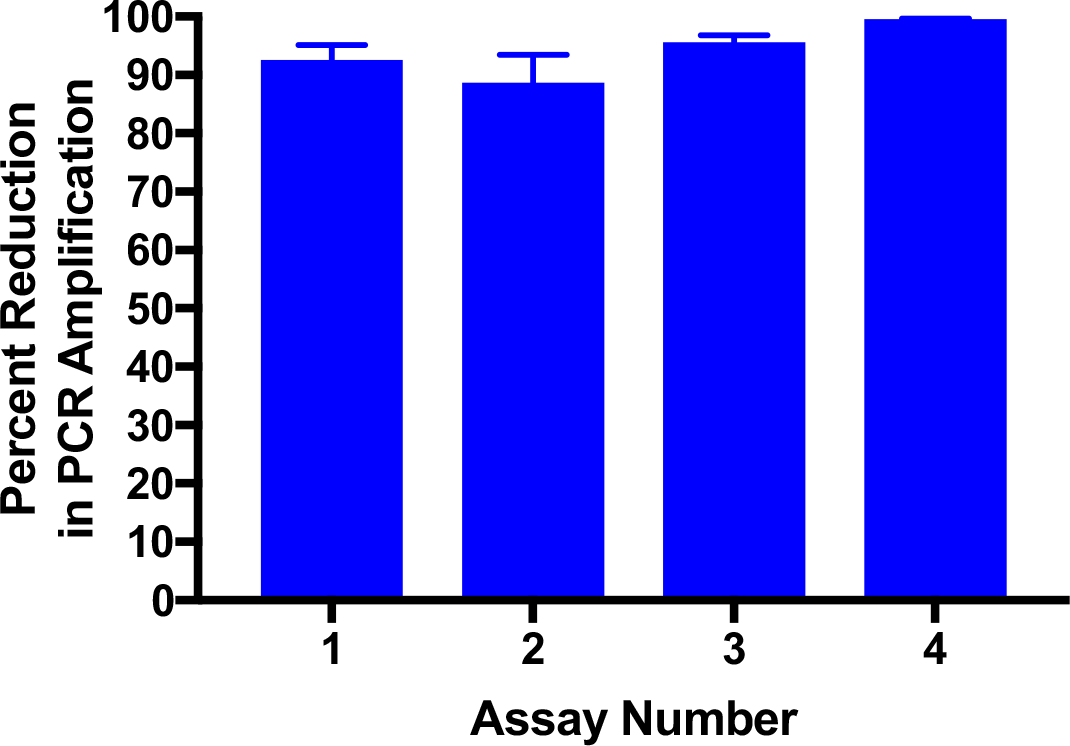
Selected assay conditions gave reproducible inhibition of PCR signal from dead cells. Heat-killed samples were treated with 50 μM of PMA and incubated in the dark for 10 minutes followed by 5 minutes of light activation. A 94.15% reduction in the PCR amplification signal was observed. Error bars represent standard deviations from five biological replicates.

### The effect of exogenous cells on the detection and PMA-mediated inhibition

Clinical samples are likely to contain cells other than *B. pertussis*, including eukaryotic cells that contain very large amounts of DNA compared to *B. pertussis* cells. Eukaryotic cells may interfere with the PMA-mediated inhibition of amplification from dead *B. pertussis* preventing distinguishing between live and dead *B. pertussis*. To test this, varying amounts of gDNA from THP-1 cells were combined with a constant amount of *B. pertussis* gDNA, and Ct values were determined and compared to samples containing *B. pertussis* only. No effect of THP-1 gDNA on detection of *B. pertussis* was observed up to an equivalent of approximately 5500 THP-1 cells per assay (Figure 6).

**Figure 6.**
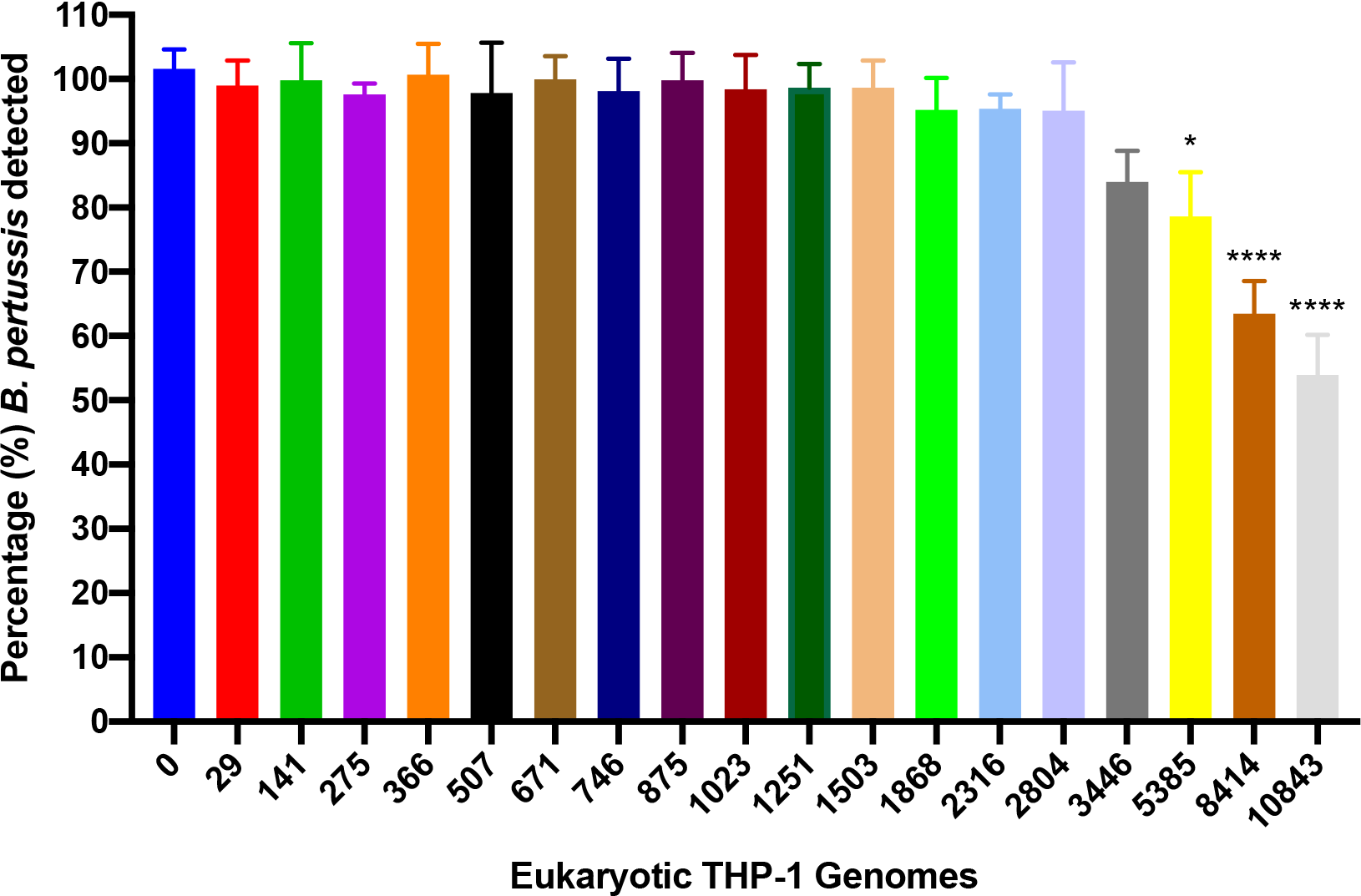
Effect of eukaryotic gDNA on detection of *B. pertussis*. Varying amounts of gDNA from THP-1 cells were combined with gDNA equivalent to 10^3^ *B. pertussis* cells. No interference in detection of *B. pertussis* was observed up to the equivalent of 3446 THP-1 cells, after which the sensitivity of detection was reduced when compared to viable *B. pertussis* detected in the presence of 0 THP-1 cells. *: p<0.05, determined by one-way ANOVA and Dunnett’s multiple comparisons test, with a single pooled variance. Error bars represent standard deviations from three biological replicates.

It was possible that the presence of other cells would interfere with the PMA-mediated inhibition of PCR signal from dead *B. pertussis*. Thus, the effect of heat-killed or live THP-1 cells on PMA-mediated inhibition of PCR amplification from heat-killed *B. pertussis* was tested. A 99.94% reduction in PCR signal was observed indicating that THP-1 cells did not prevent PMA-mediated inhibition of PCR signal from dead *B. pertussis* (Figure 7).

**Figure 7.**
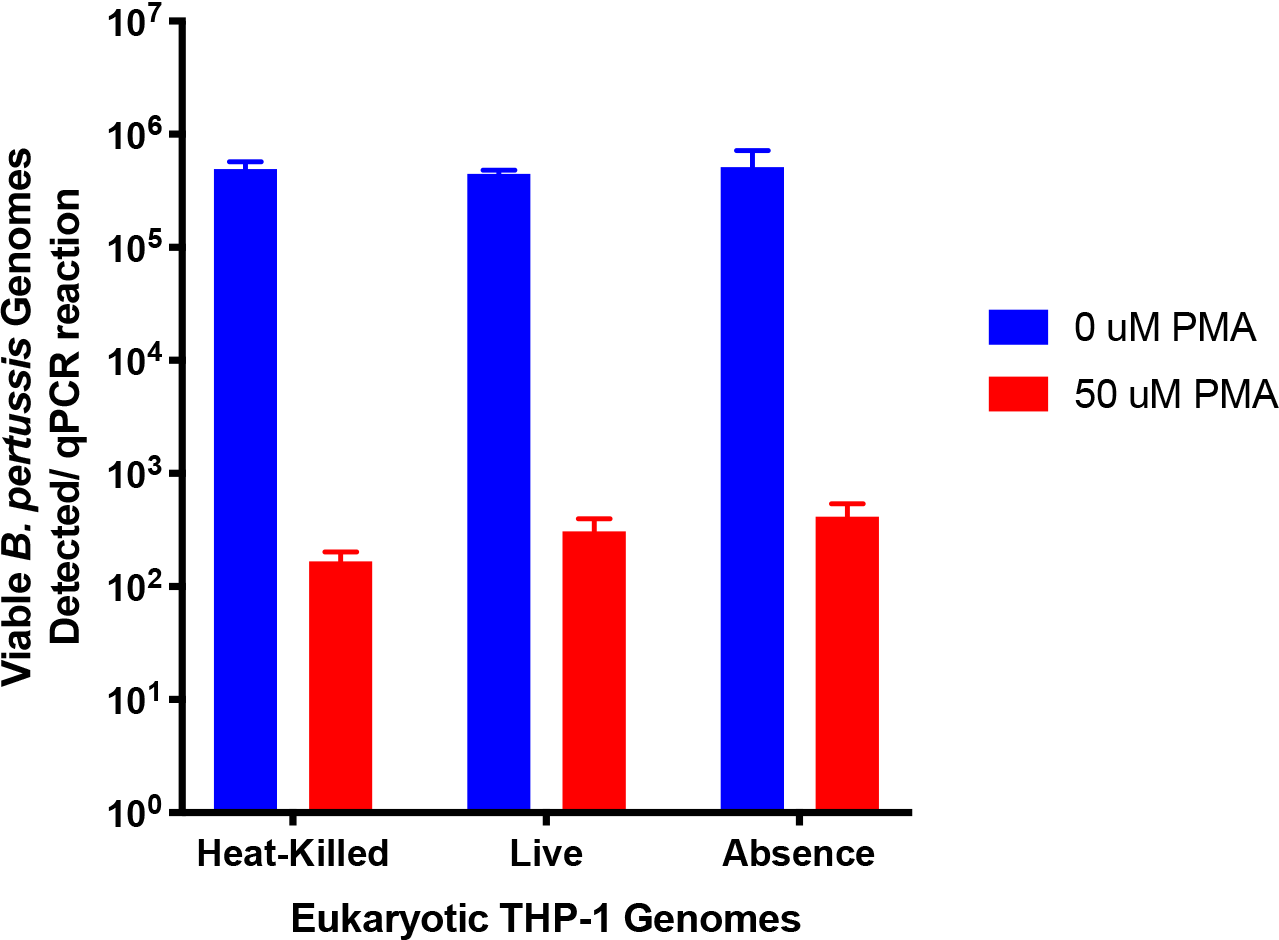
Eukaryotic THP-1 gDNA did not interfere with enumeration of *B. pertussis*. gDNA from 10^6^ heat-killed *B. pertussis* were combined with either 10^5^ heat-killed THP-1 cells, 10^5^ live THP-1 cells or a no THP-1 cell control and treated with PMA. Non PMA-treated samples were run in parallel. The presence of live or dead THP-1 cells did not interfere with the action of PMA on dead *B. pertussis* cells. Error bars represent standard deviations from three biological replicates.

To test the assay’s ability to distinguish between viable and dead *B. pertussis,* in the presence of other cells, a constant number of THP-1 cells were combined with different ratios of heat-killed and viable *B. pertussis* cells. The reduction in PCR signal was proportional to the amount of heat-killed cells in each suspension (Figure 8) demonstrating that the assay was able to distinguish viable from dead *B. pertussis*, even in the presence of human cells.

**Figure 8.**
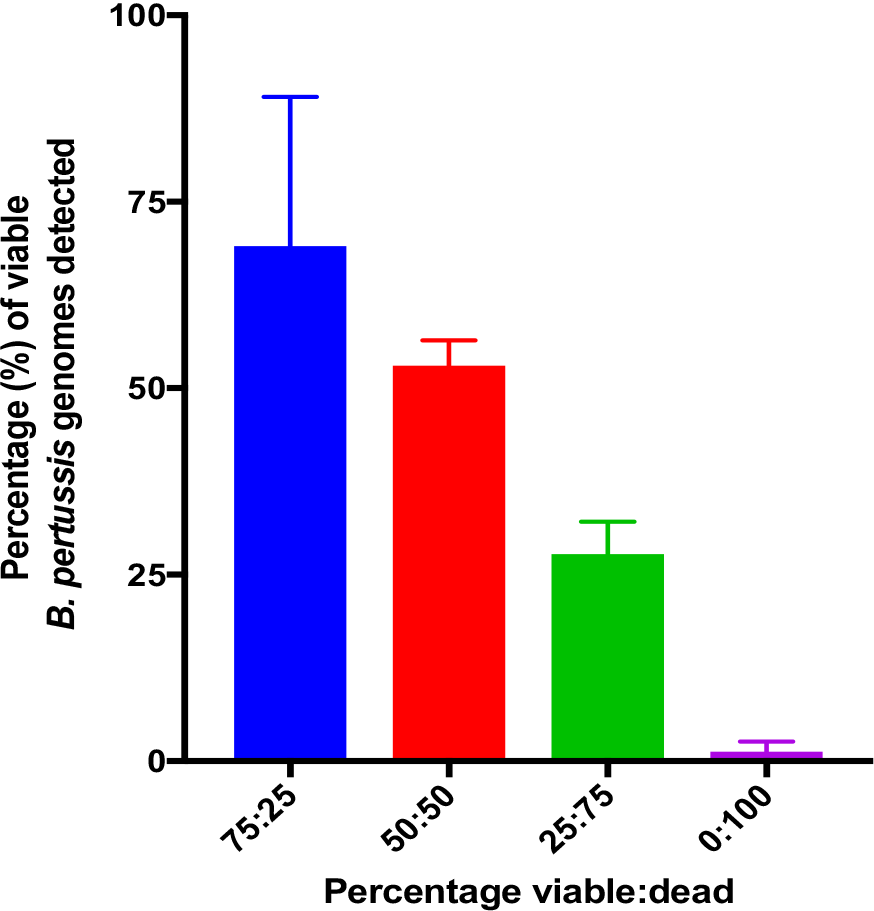
Percentage of live *B. pertussis* cells enumerated from PMA treated samples in the presence of eukaryotic cells. 100,000 THP-1 cells were combined with suspensions of different ratios of heat-killed and viable *B. pertussis* cells. The assay accurately distinguished viable from dead *B. pertussis* in each suspension. Error bars represent standard deviations from three biological replicates.

Collectively, these studies revealed that the THP-1 cells did not interfere with the PMA-mediated inhibition of PCR signal from dead *B. pertussis* or prevent the accurate enumeration of viable *B. pertussis* cells.

### Measuring the viability of *B. pertussis* during in vitro growth

During development of the assay, it was observed that PMA treatment of live *B. pertussis* suspensions used as controls consistently reduced the PCR signal compared to untreated samples. This suggested that *B. pertussis* colonies taken from plate grown cultures contains both live and dead bacteria. To investigate this, and to determine the proportion of live to dead *B. pertussis* in plate grown cultures over time, suspensions of cells were made of *B. pertussis* grown on plates for either 3, 4, 5 or 8 days. The suspensions were treated with PMA and qPCR performed. The percentage of PCR signal observed was compared to untreated controls, Figure 9. *B. pertussis* is relatively slow growing and many protocols for plate growth involve incubation for 72 hours to achieve visible colonies. However, at this point *B. pertussis* viability was only 89%. Interestingly, although colony size continued to increase between days 3 and 5, percentage viability decreased to 24%. Further incubation resulted in further loss in viability. Thus, when using plate grown *B. pertussis* in assays, suspensions will be a mixture of live and dead bacteria, and that enumeration of *B. pertussis* by plating serial dilutions of a suspension and counting the resulting CFU’s will not be a measure of the total number of cells in the suspension.

**Figure 9.**
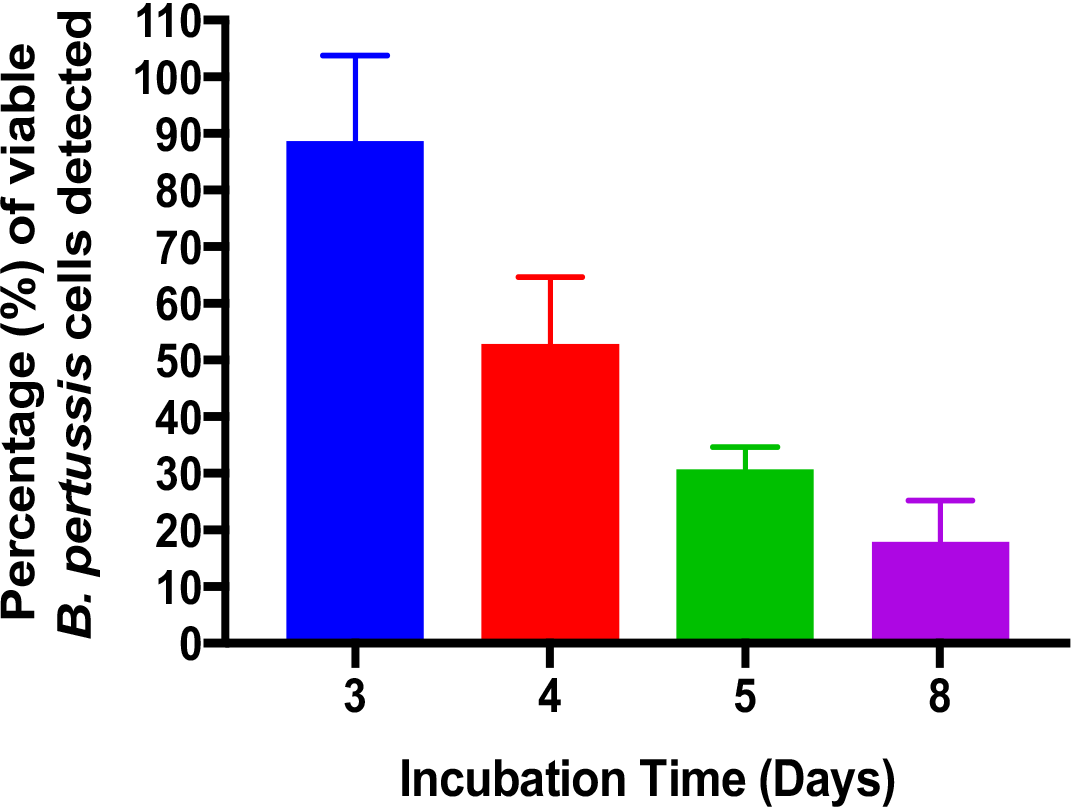
The viability of *B. pertussis* decreases during growth on agar plates. The viability of *B. pertussis* growing on agar plates was measured over time. Viability decreased as the incubation time increased with only 24% of cells being viable after 5 days of incubation. Error bars represent standard deviations from three biological replicates.

### Use of the assay to enumerate live and dead B. pertussis from human challenge model samples

The assay was developed in order to provide a method for monitoring the colonisation status of participants in a novel human challenge model of *B. pertussis* colonisation. During development of this model, a group of participants were inoculated with 10^5^ CFU of *B. pertussis* and daily samples were taken over a 14-day period to monitor colonisation (15). Samples types included nasosorption fluids, pernasal swabs, throat swabs, and nasopharyngeal washes. Samples were split and one portion was treated with PMA. *B. pertussis* were enumerated from treated and untreated samples. In addition, portions of samples were serially diluted and plated for enumeration of *B. pertussis* by traditional culture. Under these conditions, 3 out of 5 participants were determined to be colonised by culture of *B. pertussis* (data not shown). Samples obtained from volunteers on day 9 post-challenge were tested by PMA-qPCR which revealed that 4 out of 5 volunteers were deemed to carry viable *B. pertussis* by this method (Figure 10). This was observed after detectable viable *B. pertussis* were found in nasal washes and pernasal swabs. Nasal washes from Day 11 samples also had detectable viable *B. pertussis* in 2 of the 5 volunteers (Figure 10). Samples from positive volunteers contained roughly the same number of viable and dead *B. pertussis*. Interestingly, on Day 16 of sampling, two days after volunteers started azithromycin treatment to eradicate the infection, all but one volunteer was negative for detectable *B. pertussis* genomes. In this volunteer, the PMA-qPCR assay was able to detect low levels of viable and dead *B. pertussis*, with a higher proportion of dead genomes detected compared to viable genomes (Figure 10 F).

**Figure 10.**
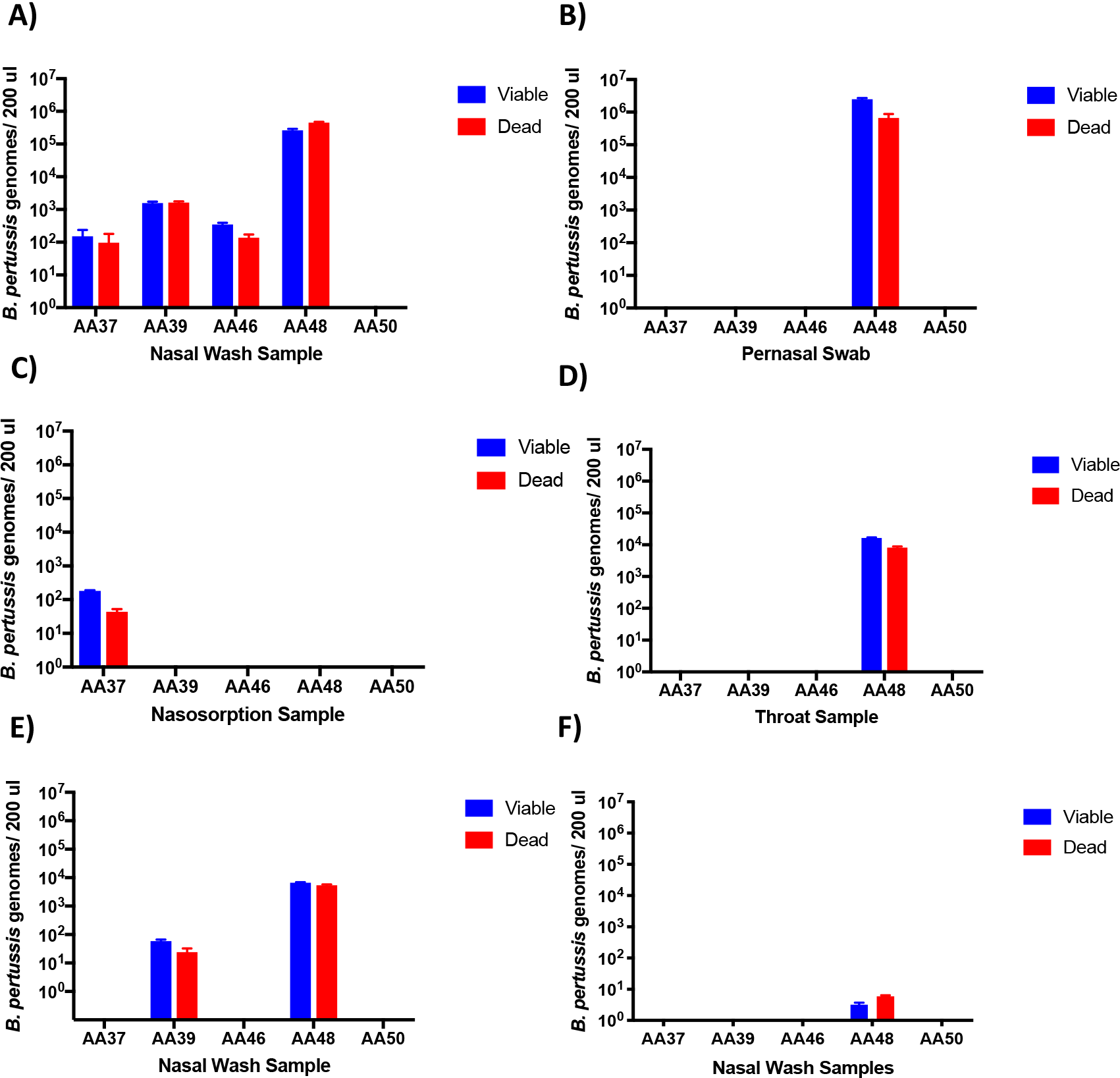
PMA-qPCR detected viable *B. pertussis* from Human Challenge Model samples within hours. Viable and dead *B. pertussis* were enumerated in samples from 5 volunteers in the Human Challenge Model, collected Day 9 (A-D), Day 11 (E) and Day 16 (F) after inoculation, from the sample type indicated. 200ul of samples were processed. Day 16 samples are taken two days after volunteers started azithromycin treatment to clear infection. Values below the lower limit of detection were considered undetectable and given a value of 0.

## Discussion

Ordinarily, the detection and quantification of viable *B. pertussis* is achieved through culture on laboratory agar. However, the relatively slow growth rate of *B. pertussis* means that the growth of countable colonies can take between 72 – 120 hours. The development of a human challenge model for *B. pertussis* as part of the PERISCOPE project requires that enumeration of viable *B. pertussis* be achieved in a much shorter time than this, in order to be able follow colonisation closely. In addition, simple enumeration of viable bacteria within a sample doesn’t provide the complete picture. In many scenarios, such as measuring bacterial load in an infection model, it is of great interest to know the total bacterial number as understanding the dynamics of bacterial growth that involves both cell division and cell death is very important. Thus, while traditional qPCR provides a faster detection method for *B. pertussis* than culture, the modification of a qPCR assay with the introduction of PMA treatment of samples reported here enables both fast detection of *B. pertussis* and the ability to distinguish viable from dead cells.

Here, we demonstrate that PMA inhibits PCR-mediated amplification from dead *B. pertussis* and that inhibition of signal from dead cells occurs even in the presence of high numbers of eukaryotic cells. This may be important for the detection of *B. pertussis* from complex samples that contain a mix of cell types as seen in the human challenge model. Samples obtained from volunteers that were identified as positive for *B. pertussis* by culture, were also detected in our initial test of the PMA-qPCR assay. The same volunteers were identified as being negative for *B. pertussis* by both qPCR and culture, with the exception of a single sample that had low levels of *B. pertussis* identified only by qPCR. Interestingly, PMA-qPCR detected approximately equal numbers of viable and dead *B. pertussis,* demonstrating its use to enumerate total bacteria rather than only viable. The full results of the human challenge model are published elsewhere (15). Here we demonstrate that the PMA-qPCR assay allowed for a determination of colonisation status within hours of obtaining the samples compared to days when using culture.

The utility of the PMA-qPCR assay has been shown in the human challenge model, but this assay has other uses. For example, in diagnostic laboratories, where ascertaining if *B. pertussis* is viable or dead will facilitate whether to pursue culture as a means to obtaining a live culture for characterisation. It is also of use in a range of research and industrial settings enabling investigation of the dynamics of *B. pertussis* growth by determining both cell division and cell death.

## Acknowledgments

This work was supported by funding to the PERISCOPE Consortium. PERISCOPE has received funding from the Innovative Medicines Initiative 2 Joint Undertaking under grant agreement No 115910. This Joint Undertaking receives support from the European Union’s Horizon 2020 research and innovation programme and EFPIA and BMGF.

